# MpsLDA-ProSVM: predicting multi-label protein subcellular localization by wMLDAe dimensionality reduction and ProSVM classifier

**DOI:** 10.1101/2020.04.19.049478

**Authors:** Qi Zhang, Shan Li, Bin Yu, Yang Li, Yandan Zhang, Qin Ma, Yusen Zhang

## Abstract

Proteins play a significant part in life processes such as cell growth, development, and reproduction. Exploring protein subcellular localization (SCL) is a direct way to better understand the function of proteins in cells. Studies have found that more and more proteins belong to multiple subcellular locations, and these proteins are called multi-label proteins. They not only play a key role in cell life activities, but also play an indispensable role in medicine and drug development. This article first presents a new prediction model, MpsLDA-ProSVM, to predict the SCL of multi-label proteins. Firstly, the physical and chemical information, evolution information, sequence information and annotation information of protein sequences are fused. Then, for the first time, use a weighted multi-label linear discriminant analysis framework based on entropy weight form (wMLDAe) to refine and purify features, reduce the difficulty of learning. Finally, input the optimal feature subset into the multi-label learning with label-specific features (LIFT) and multi-label k-nearest neighbor (ML-KNN) algorithms to obtain a synthetic ranking of relevant labels, and then use Prediction and Relevance Ordering based SVM (ProSVM) classifier to predict the SCLs. This method can rank and classify related tags at the same time, which greatly improves the efficiency of the model. Tested by jackknife method, the overall actual accuracy (OAA) on virus, plant, Gram-positive bacteria and Gram-negative bacteria datasets are 98.06%, 98.97%, 99.81% and 98.49%, which are 0.56%-9.16%, 5.37%-30.87%, 3.51%-6.91% and 3.99%-8.59% higher than other advanced methods respectively. The source codes and datasets are available at https://github.com/QUST-AIBBDRC/MpsLDA-ProSVM/.

## 1. Introduction

The nature of proteome research refers to studying the characteristics of proteins in a large range, including protein-protein interactions [1, 2], post-translational modifications [3], protein expression levels [4], etc. From this, we can get a comprehensive understanding of cell metabolism, disease occurrence and other processes at the protein level. However, proteins exert their characteristics only in the corresponding subcellular location [5], so determining the protein subcellular location is of great significance for understanding the protein characteristics and functional diversity [6], which is considered to lay the foundation for protein design and synthesis. In recent years, researchers have put forward a number of forecasting methods to predict SCLs, but most of the predictors can only process unit proteins [7]. However, experiments have found that some proteins can appear in two or more different Subcellular position, which are called multi-label proteins [8]. For example, acyl-CoA synthetase is found in both the endoplasmic reticulum and mitochondrial outer membrane [9]; Antioxidant defense proteins are present in mitochondria, peroxisome and cytosol [10].These multi-label proteins play a critical role in physical transportation, information transferring, energy conversion and other life activities of cells [11]. Therefore, it is urgent to explore the SCL of multi-label proteins.

The first task of SCL prediction of multi-label proteins is feature extraction. At present, the commonly used features include evolutionary information, physical and chemical properties, sequence information and structure information. For example, in order to explore the function of viral proteins, Thakur et al. [12] used three different feature extraction algorithms dipeptide composition (DC), amino acid composition (AAC), physicochemical properties (PHY) and their hybrids to develop a server “MSLVP” for predicting SCLs. Zhou et al. [13] used position-specific scoring matrix (PSSM), amino acid composition (AAC), conserved domain database (CDD) and gene ontology (GO) annotation to extract and fuse the features of human proteins and propose a new method Hum-mPLoc 3.0 based on amino acid sequence. Cheng et al. [14] fused GO information and pseudo amino acid composition (PseAAC) to extract features and predict the multiple subcellular locations from animal dataset.

However, single feature extraction method cannot fully characterize the feature information in the sequences, so researchers take the way of feature fusion to characterize the proteins multidimensionally. But the high-dimensional data after fusion contains a lot of irrelevant redundancy, which will seriously mislead the prediction. Dimensionality reduction can solve such problems, remove irrelevant information, and improve the running speed of the model. At present, Li et al. [15] constructed a robust multi-label kernelized fuzzy rough set model named RMFRS. The model combined the kernel information of sample space and label space, and combined multi-core learning with fuzzy rough set to complete feature selection. According to information theory, Zhang et al. [16] put forward a feature selection method named multi-label feature selection based on label redundancy (LRFS), which used conditional mutual information between each label and candidate features, and combined the feature redundant terms to determine the final feature vector. Chen et al. [17] put forward a new method of extended adaptive least absolute shrinkage and selection operator (EALasso). This method preserves the oracle properties of identifying the correct subset model and has the optimal estimation accuracy. The convergence of the algorithm was achieved by using the iterative optimization algorithm.

Machine learning algorithm such as classifiers acts as the last step for SCL prediction of multi-label proteins. Recently, Wang et al. [18] used the algorithm of integrating multiple classifier chains (ECC) for prediction on the multi-label protein datasets of Gram-negative bacteria and Gram-positive bacteria. The overall prediction accuracy (OLA) and the overall actual accuracy (OAA) about these two datasets were 94.1%, 92.4% and 94.4%, 94.0%, respectively. Wan et al. [19] built an integrated multi-label classifier HPSLPred, which was proved to have the highest accuracy and average precision of 75.89% on human datasets. Javed and Hayat [8] used ranking support vector machine and multi-label k-nearest neighbor classifiers for prediction about virus and Gram-positive bacteria datasets. The overall accuracy on the two datasets were 80.46% and 85.54%, respectively.

Although existing prediction methods have achieved good prediction performance, in order to further improve the prediction performance, it is necessary to build a prediction model by combining multiple information, processing high-dimensional data, and selecting the appropriate classifier. Inspired by this, this paper proposes a new algorithm called MpsLDA-ProSVM to predict the SCL of multi-label proteins. Firstly, four feature coding methods, PseAAC, CT, PsePSSM and GO, are used to draw the feature information from protein sequences, and the information is feature-fused. Secondly, for the first time, weighted multi-label linear discriminant analysis framework based on entropy weight form (wMLDAe) is used to remove redundancy and get the best feature vectors. At last, these feature vectors are first input to the ML-KNN and LIFT algorithms to obtain the label score of each instance, so as to get the classification and ranking of the relevant labels, and then predict the multi-label proteins through the ProSVM classifier to determine the final prediction model. In this paper, two training datasets (virus and plant datasets) and two test datasets (Gram-positive bacteria and Gram-negative bacteria datasets) are used for evaluation by jackknife method, and the overall actual accuracy (OAA) on virus, plant, Gram-positive bacteria and Gram-negative bacteria datasets are 98.06%, 98.97%, 99.81% and 98.49%, respectively, and the overall location accuracy (OLA) are 98.41%, 99.33%, 99.81% and 99.04% respectively. By experiment, our proposed MpsLDA-ProSVM method can helpfully improve the performance of multi-label protein SCL prediction.

## 2. Materials and methods

### 2.1. Datasets

In this study, datasets of virus proteins and plant proteins are selected as training set to construct the model, and datasets of Gram-negative bacteria and Gram-positive bacteria are selected as test set to examine the feasibility of the model. There are 207 different protein sequences in the virus dataset [20], which are located in 6 subcellular positions, among which 165 belong to one position, 39 belong to two positions and 3 belong to three positions. The plant dataset [21] has 978 different protein sequences, which are located in 12 subcellular positions. Among them, 904 belong to one position, 71 belong to two positions, and 3 belong to three positions. The breakdowns of these two training sets are listed in Fig. 1 and Fig. 2.

**Fig. 1.**
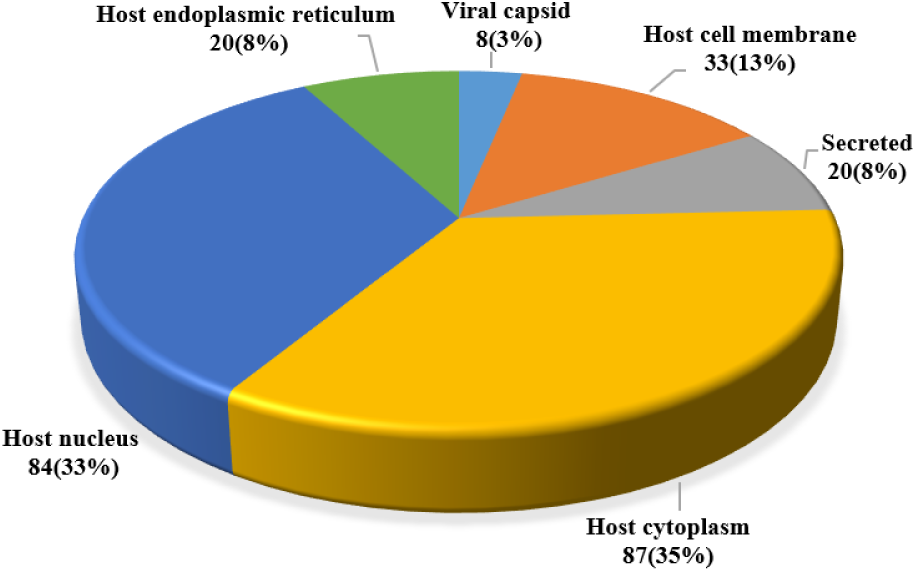
The distribution of virus dataset in different subcellular locations. The number of proteins represents the number of “location proteins” in the corresponding subcellular location. There are 252 protein sequences in six subcellular locations of the virus dataset, and the homology of any one protein with other protein sequences in the same subcellular location is limited to 25%.

**Fig. 2.**
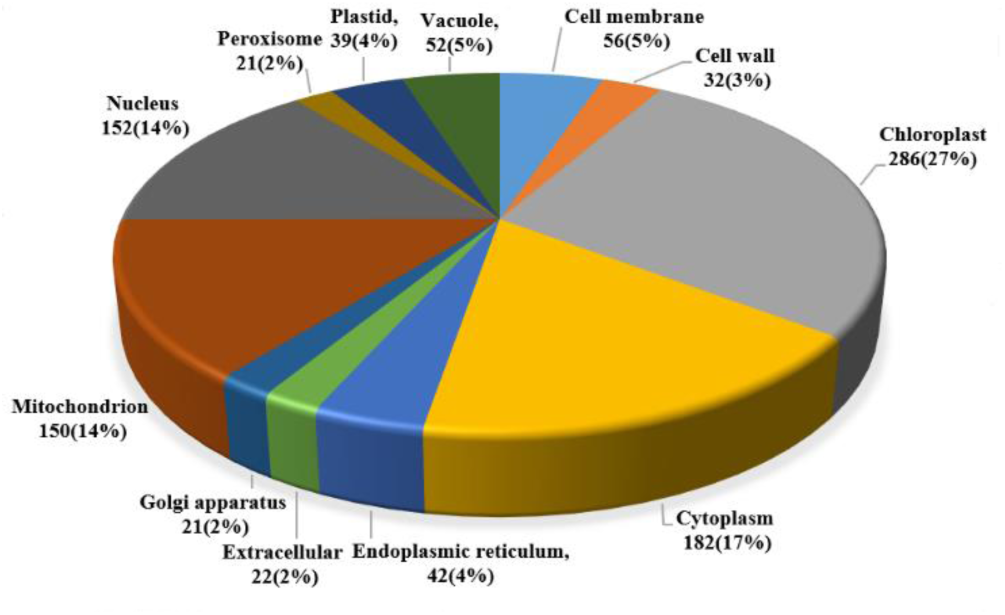
The distribution of plant dataset in different subcellular locations. The number of proteins represents the number of “location proteins” in the corresponding subcellular location. There are 1055 protein sequences in twelve subcellular locations of the plant dataset, and the homology of any one protein with other protein sequences in the same subcellular location is limited to 25%.

The Gram-positive bacteria [22] contains 519 different protein sequences, which are located in 4 subcellular positions, among which 515 belong to one position and 4 belong to two positions. The Gram-negative bacteria [23] contains 1392 distinct protein sequences, which are located in 8 subcellular locations, of which 1328 belong to one location and 64 belong to two locations. Showing the specific classification of protein sequences about two test sets are in Supplementary Table S1 and S2. The homology of any one protein with other protein sequences in the same subcellular location is limited to below 25%.

### 2.2. Feature extraction

In the article, we use pseudo position-specific scoring matrix, GO, pseudo amino acid composition and conjoint triad to draw the information from sequence and construct the initial feature vector.

#### 2.2.1. Pseudo Amino Acid Composition

The pseudo amino acid composition (PseAAC) [7] model put forward by Chou [24] can not only extract the location information of protein sequence, but also reflect the essential composition information. According to the characteristics of 20 amino acids, each amino acid sequence is normalized to (20 + *λ*) -dimensional vector, in which the first 20 dimensions represent amino acid composition information, while the last *λ* dimensions represent pseudo amino acid composition information. At this point, the sequence is converted to the following feature vector:

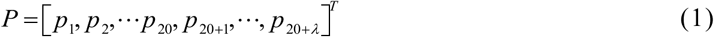

Among them,

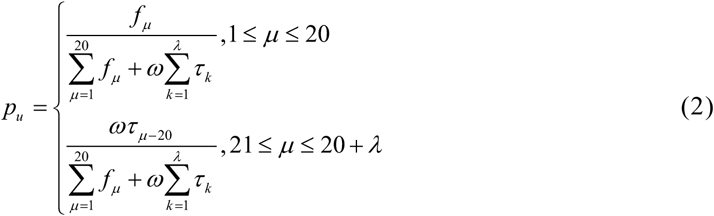

where *ω* represents the weighting factor, the value is 0.05 [25], and refers to the order information between the amino acids in the protein sequence. In this study, the value of *λ* is determined by the prediction accuracy, which is the optimal parameter value.

#### 2.2.2. Pseudo position-specific scoring matrix

Pseudo position-specific scoring matrix (PsePSSM) [26] draws evolution information from sequences with this method following two steps. First, in the NR database, align the protein sequences using the PSI-BLAST [27] program to obtain a position-specific scoring matrix (PSSM) [28]. For protein sequences of length *L*, the PSSM algorithm obtains *L* × 20 -dimensional feature vectors, and the results are as follows:

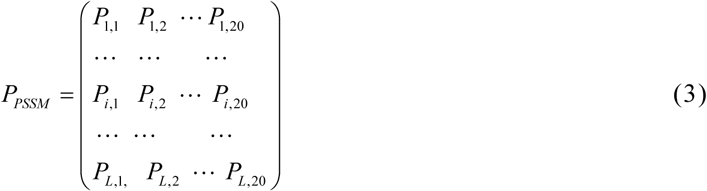

where the row indicates the corresponding amino acid position in the sequence, and the 20 amino acid types that can be mutated are listed.

Secondly, the PSSM matrix is standardized, and then use PsePSSM algorithm to generate (20 + 20×*ξ*) -dimensional eigenvectors for each protein sequence in order to unify the dimensions of the matrix. The feature extraction results are as follows:

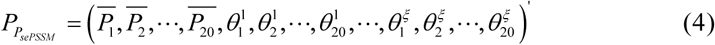

where 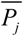 represents the component of amino acid *j* in PSSM. 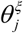 is the *ξ* -order related factor for the *j* -type amino acid. Similarly, the value of *ξ* depends on the prediction accuracy, which is the optimal feature extraction parameter.

#### 2.2.3. Conjoint triad

In order to draw sequence information from proteins, Shen et al. [29] proposed conjoint triad (CT) [30, 31] algorithm based on the dipole and volume of amino acid side chains. First, according to the dipole and volume of the amino acid side chain, the 20 amino acids are divided into seven categories. Secondly, every three of the protein sequences are grouped to form a triplet. By analogy, each sequence can generate 7×7× 7=343 triplets. Calculate the frequency of each triplet to form a 343-dimensional feature vector. The formula is as follows:

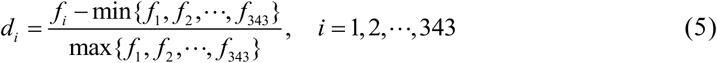

#### 2.2.4. Gene ontology

Gene ontology is a new expression method based on GO database, which is used to draw annotation information from protein sequence. The GO model uses GO ID to annotate the characteristics of protein sequences. It annotates genes and proteins based on three aspects: biological processes, molecular functions, and cell composition. Use BLASTP to search for homologous proteins in the SWISS-PROT database and remove sequences with similarities ≥ 60%. For the remaining protein sequences *S*_*i*_, the GO information [32, 33] of the corresponding protein is determined in the GO database according to the index number in the sequence. Its manifestation is:

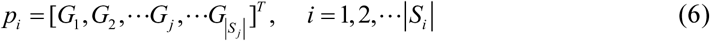

where |*S*_*i*_| is the size of the *S*_*i*_. If the GO number exists, *G*_*j*_ = 1 ; otherwise, *G*_*j*_ = 0.

### 2.3. Weighted multi-label linear discriminant analysis framework based on entropy weight form

Linear discriminant analysis is one of the most widely used feature selection methods. For multi-label data, Xu [34] proposed a weighted multi-label linear discriminant analysis framework based on binary and related weight forms. Assuming *l* multi-label training instances and *q* class label sets are given, define *W* as *l* × *q* -dimensional non-negative weight matrix, that is:

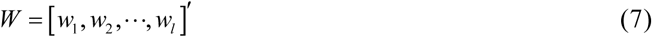

where *w*_*i*_ = [*w*_*i*1_, …, *w*_*ik*_, …, *w*_*iq*_]′, *i* < *l* represents the *i*-*th* instance, define the between-class, within-class and total scatter matrices as:

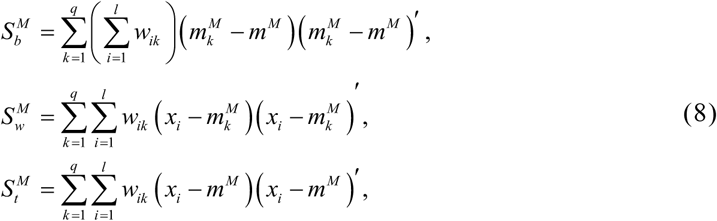

where *m*^*M*^ is the total mean vector and 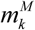 is the mean vector of the *k* -*th* class. Chen [35] put forward an entropy-based weight form and is introduced into the feature selection process of this section. Entropy is a metric to measure the uncertainty of random variables [36]. To the *i*-*th* instance, the entropy can be defined as:

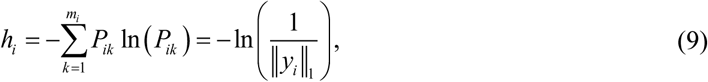

Let *w*_*ik*_ indicate the probability that the *i*-*th* instance belongs to the label of the *k* -*th* class, which can be estimated by formula (10):

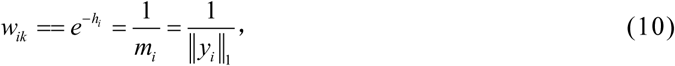

This shows that the probability of the relevant labels is equal to the inverse of the number of labels. At this time, the sum of the weights of each instance is 1, and the total scatter matrix is:

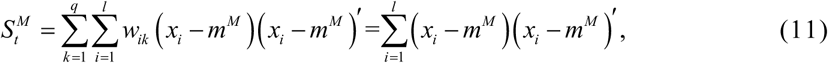

The results of the formula (11) are defined in the same way as those of the single-label LDA algorithm [37]. In the wMLDA algorithm, the projection matrix P is obtained by solving the generalized eigenvalue problem, that is:

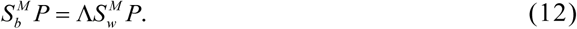

where Λ is composed of *C* non-negative eigenvalues, that is *λ*_1_ ≥ *λ*_2_ ≥ … ≥ *λ*_*C*_ ≥ 0. A wMLDA having the form of the above entropy weight is added with a suffix ‘e’, which is called wMLDAe. In this paper, the algorithm is used to process the fused vector, taking full account of the label information, avoiding over-fitting of the model, and improving the running speed.

### 2.4. Prediction and Relevance Ordering based SVM

Assume that a set of *n* training instances 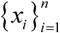 and *T* label *L* = {*l*_1_, *l*_2_,…, *l*_*T*_} are given. In order to consider the comprehensive ranking of relevant labels, we first use the ML-KNN [38] and LIFT [39] algorithms to obtain the label score of each training instance, denoted as *g*(*x*_*i*_) = [*g*_1_ (*x*_*i*_), *g*_2_ (*x*_*i*_),…, *g*_*T*_ (*x*_*i*_)]. Each label *l*_*t*_, *t* ∈{1, 2,…,*T*} is assigned a real value *g*_*t*_ (*x*_*i*_), and the labels are sorted according to the real values to obtain the corresponding set *R*_*i*_ ∈ *L* of the ranking of relevant labels. *φ*_*x*_ (*a*) is used to represent the label index set that is less relevant than *l*_*a*_. Both ML-KNN and LIFT algorithm can be used for nonlinear classification, and have solid theoretical basis, low training time complexity and high stability of prediction algorithm. The purpose of Prediction and Relevance Ordering based SVM (ProSVM) [40] is to optimize the PRO Loss function, which is non-convex and difficult to optimize. Therefore, we consider optimizing its equivalent loss function as follows:

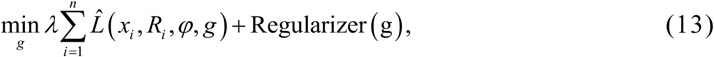

where

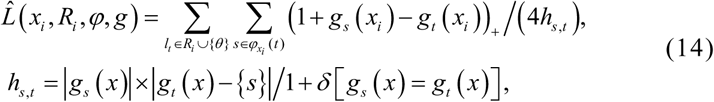

Regularizer (*g*) is a regularizer of *g, λ* is a parameter measuring functional complexity of *g* and the equivalent loss function.

Suppose *g*′*s* are linear models with 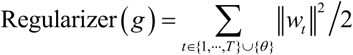 and 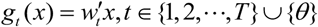, in which *θ* is the threshold. Let *G* be the training set, 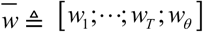, and (13) can be transformed into SVM-type in general form:

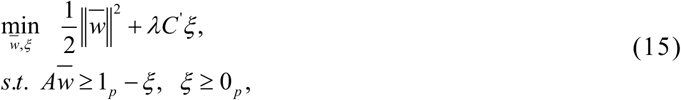

where 1_*p*_ is the *p* ×1 -dimensional one matrix and *p* is the total number of constraints. The vector *C* represents the weight of the loss, and the matrix *A* represents the constraint value between samples. *ξ* is determined by the value of 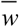.

The Alternating Direction Method of Multipliers (ADMM) [41] is a widely used optimization method for constraint problems in machine learning. The idea of decomposition and coordination is used to find the answer of the primitive question by solving its sub problems. Its basic methodology is to use the augmented LaGrange method to solve the optimization problem with equality constraints. Decompose the training set *G* into *N* disjoint subsets. Fusion the objective function and constraints through a dual variable to form an augmented LaGrange function. According to the ADMM solution algorithm, the specific process is as follows:

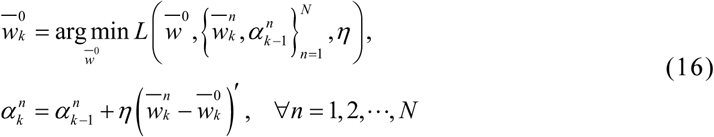

Through the above method, it is equivalent to solve the following problem:

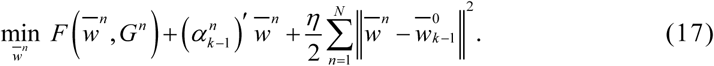

This is a quadratic programming (QP) issue and Eq. (17) is similar to the standard SVM problem. At this time, the multi-label problem is converted into multiple binary classification problems (each label corresponds to a binary classification), which can be solved by the latest SVM solver LIBLINEAR [42] to obtain the score of each instance, and classify the instances by the threshold.

### 2.5. Performance evaluation

According to Chou’s five-step rule [43], choosing appropriate indicators to judge the quality of a prediction system is an important steps to build a prediction model. At present, the commonly used inspection methods include re-substitution, independent test, k-fold cross-validation and jackknife. Among them, jackknife method used in this paper is regarded as one of the most rigorous inspection methods [26]. For the jackknife test, specifically, any one of the N protein sequences is selected as testing, and the other N-1 sequences are used as training, which is repeated for N times until all protein sequences are tested.

So as to more intuitively assess the prediction performance about the model, we use eight measures including Hamming Loss (HL), Average Precision (AP), Ranking Loss (RL), F1, PRO Loss (PL), Coverage (CV), overall location accuracy (OLA) and overall actual accuracy (OAA) [40, 44, 45]. Suppose a given multi-label training set 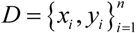 and *T* learning functions *F* = {*f*_1_, *f*_2_,…, *f*_*T*_}, where *y*_*i*_ ∈{1, − 1}^*T*^ is the real label of example *x*_*i*_, and *H* = {*h*_1_, *h*_2_,…, *h*_*T*_} represents *T* multi-label classifiers. 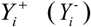 represents the true relevant (irrelevant) labels set of instance *x*_*i*_, and 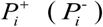 represents the predicted relevant (irrelevant) labels set of instance *x*_*i*_. *rank*_*F*_ (*x*_*i*_, *j*) represents the ranking of *j* label in the label ranking list. The specific definitions of the indicators are as follows:

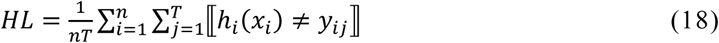

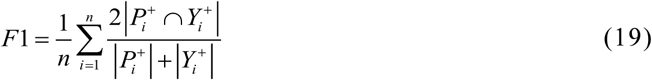

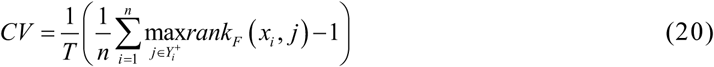

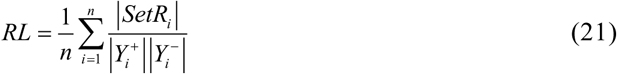

where 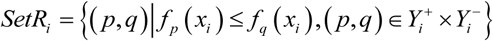.

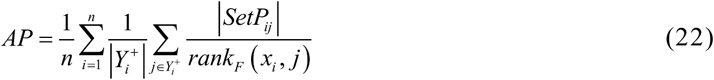

where 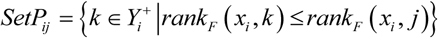.

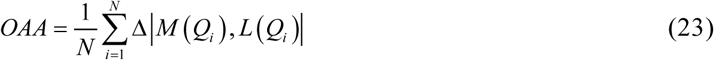

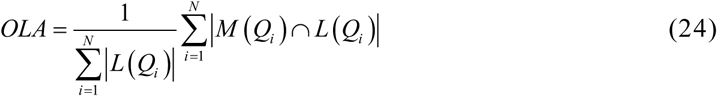

where *M* (*Q*_*i*_), *L* (*Q*_*i*_) represents the predicted label subset and the real label subset respectively, and

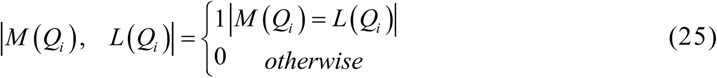

According to the definition in 2.4, this paper uses the PRO Loss evaluation index, which refers to the ranking of relevant labels and the predicted value of labels. Specifically defined as:

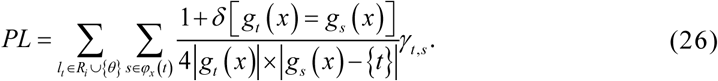

where *γ* _*t, s*_ is the modified 0-1 error. Specifically, *γ*_*t, s*_ = 1 if *g*_*t*_ (*x*) < *g*_*s*_ (*x*), *γ*_*t, s*_ = 1/2 if *g*_*t*_ (*x*) = *g*_*s*_ (*x*), otherwise the value is 0.

For PL, HL, CV and RL, the smaller the predicted value is, the better the classification performance. For AP, F1, OAA, and OLA, the larger the predicted value is, the better the classification performance.

### 2.6. Illustration of the MpsLDA-ProSVM workflow

For convenience, in this article, we put forward a multi-label protein SCL prediction method, MpsLDA-ProSVM, and the pipeline is shown in Fig. 3. The experiment environment is: windows Server 2012R2 Intel (R) Xeon (TM) CPU E5-2650 @ 2.30GHz 2.30GHz with 32.0GB of RAM, MATLAB2014a.

**Fig. 3.**
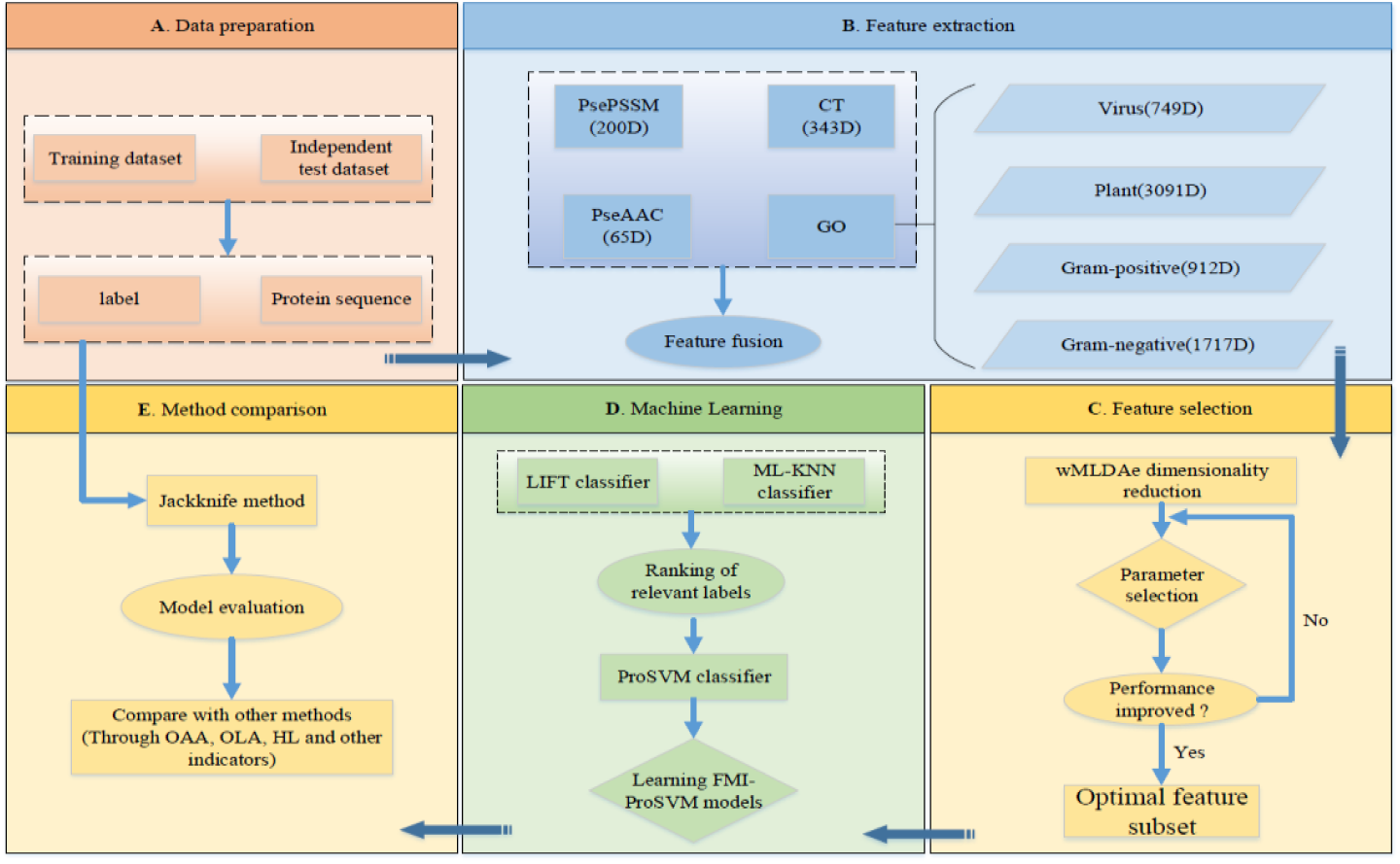
The prediction pipeline of MpsLDA-ProSVM. (A) The protein sequences of the training and testing sets and their label vectors. (B) PsePSSM, CT, PseAAC, GO are used for feature extraction and fusion. (C) wMLDAe algorithm is used to reduce dimensions and select the optimal feature subset. (D) First, we use ML-KNN and LIFT algorithm to get the comprehensive ranking of relevant labels, then input it into ProSVM classifier for prediction, and construct the MpsLDA-ProSVM model. (E) Tested by the jackknife method and compared with other methods in the paper to assess the performance of the model.

The specific steps of the MpsLDA-ProSVM method are:

***Step 1:*** Obtain the protein sequences of virus and plant datasets and make the true class labels of the corresponding sequences according to the classification.

***Step 2:*** Extract the protein sequences to get the feature information by using PsePSSM, CT, PseAAC and GO. The parameters of feature extraction are determined by inputting feature information into ProSVM classifier.

***Step 3:*** Combine the vectors from four feature extraction methods, and use wMLDAe algorithm to select the best feature subset.

***Step 4:*** Input the features selected in ***Step 3*** to the ML-KNN and LIFT classifiers to obtain the classification and ranking of relevant labels, and then use the ProSVM classifier test to perform multi-label protein SCL prediction by the jackknife, leading to the MpsLDA-ProSVM model.

***Step 5:*** Use the Gram-positive bacteria and Gram-negative bacteria datasets as independent test sets, and HL, OAA, OLA and other measures as final evaluation indicators to assess the performance of the MpsLDA-ProSVM model.

## 3. Results and discussion

### 3.1. Selection of optimal parameters λ and ξ

In order to build the optimal protein SCL prediction model and extract the optimal feature vectors from the protein sequences, the parameters of the model need to be continuously adjusted. Specially, the settings of the built-in parameters *λ* and *ξ* in the algorithm will directly affect the model performance for virus and plant datasets. To determine the values of the parameters *λ* and *ξ*, we set the *ξ* value to 1 to 10 with an interval of 1. Set the *λ* value at 5 intervals. Because the shortest protein sequence length in the independent test set is 50, the value of parameter *λ* ranges from 5 to 49. The results of feature extraction under different parameters are input into the ProSVM classifier, tested by jackknife method, and PL is selected as an evaluation index to determine the final parameters. The change of PL value of virus and plant datasets by selecting different parameter values are shown in Fig. 4. The specific results of other evaluation indexes are shown in Supplementary Table S3 and S4.

**Fig. 4.**
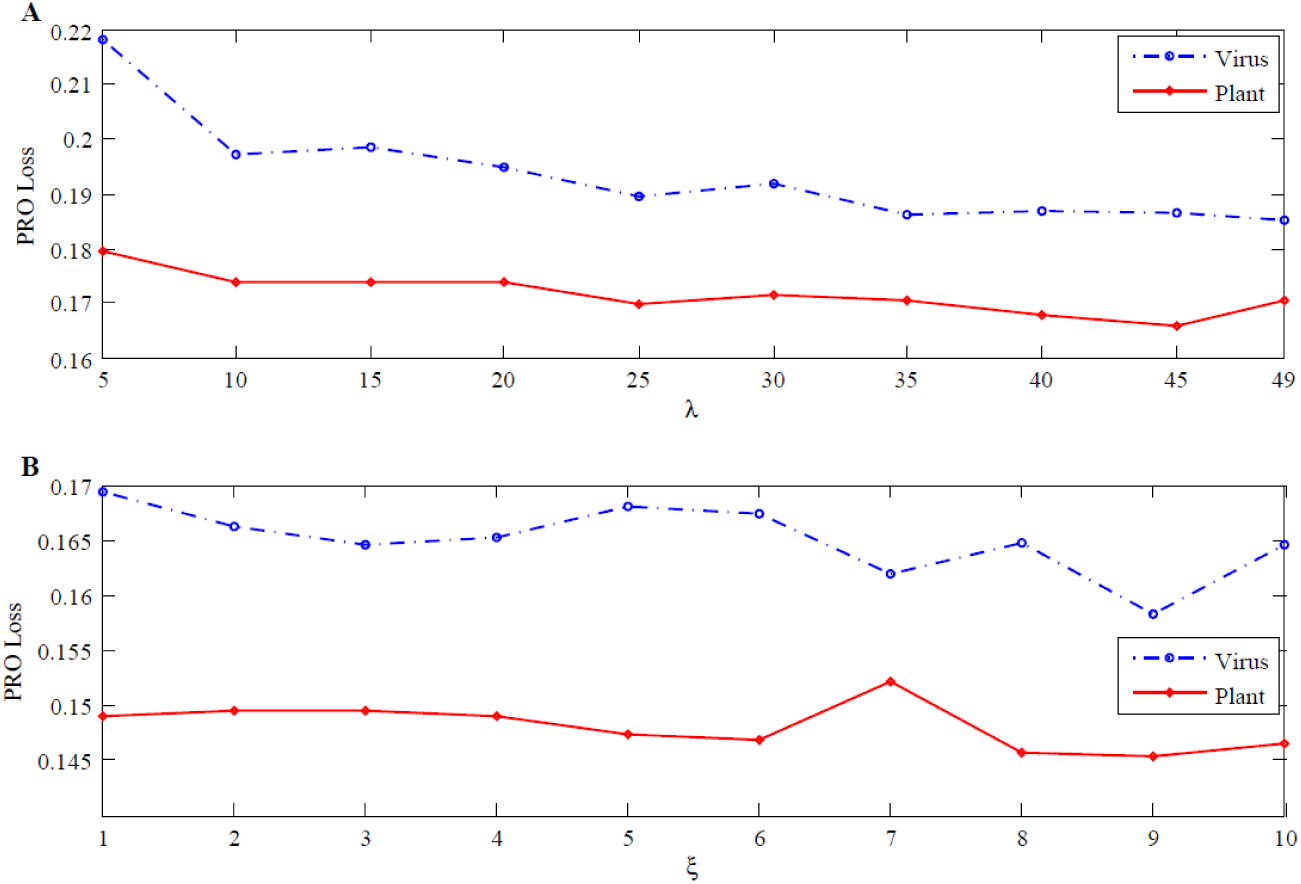
The effect of different parameters on PL value of virus and plant datasets. (A) The PL value obtained using the PseAAC algorithm; (B) The PL value obtained using the PsePSSM algorithm.

It can be seen from Fig. 4 that the prediction result of the classifier varies with different parameter values. The smaller the PL value, the better the performance of the model. For the value of parameter *λ, λ* =45 is selected under comprehensive consideration. At this time, when using PseAAC for feature extraction, a total of (20+*λ*=65) -dimensional feature vectors are generated. For the value of parameter *ξ*, when using PsePSSM algorithm for feature extraction, select *ξ*=9. At this time, a protein sequence generates a (20+20×*ξ* =200) -dimensional feature vector.

### 3.2. Effect of feature extraction algorithm on results

The PseAAC and PsePSSM algorithm are used to draw the physical and chemical information and evolution information from sequence. CT algorithm and GO annotation are used to draw the sequence information and annotation information. However, using only a single feature coding method cannot fully extract useful information about protein sequences, so researchers have fused the feature vectors extracted from various kinds of information together. To deal with the “dimensional disaster” after fusion, wMLDAe is used to reduce the number of dimensions generated after fusion. To verify the superiority of feature fusion, we use ProSVM classifier to predict the information after single-feature and multi-feature fusion. The jackknife method is used for testing, and PL, HL, RL, and F1 are used as evaluation indicators to assess the performance. The prediction results of virus and plant datasets using four single feature extraction algorithms (GO, PsePSSM, CT, PseAAC) and a fusion method using wMLDAe dimensionality reduction (PseAAC+PsePSSM+CT+GO(wMLDAe)) are shown in Table 1.

**Table 1.** Comparison of the results of different feature extraction algorithms on virus and plant datasets.

| Datasets | Algorithms | PL | HL | RL | AP | CV | F1 |
| --- | --- | --- | --- | --- | --- | --- | --- |
| Virus | PseAAC | 0.1864 | 0.2326 | 0.1741 | 0.7270 | 0.2842 | 0.5146 |
|  | PsePSSM | 0.1583 | 0.2512 | 0.1630 | 0.7574 | 0.2439 | 0.5607 |
|  | CT | 0.2504 | 0.2271 | 0.5750 | 0.6235 | 0.3712 | 0.4166 |
|  | GO | 0.0413 | 0.0435 | 0.0976 | 0.9246 | 0.4734 | 0.8969 |
|  | ALL | **0.0133** | 0.0064 | 0.0144 | 0.9961 | 0.0466 | 0.9885 |
| Plant | PseAAC | 0.1660 | 0.2292 | 0.1717 | 0.5815 | 0.0841 | 0.3861 |
|  | PsePSSM | 0.1454 | 0.2162 | 0.1559 | 0.6139 | 0.0859 | 0.4159 |
|  | CT | 0.2225 | 0.2049 | 0.5426 | 0.4302 | 0.2821 | 0.3235 |
|  | GO | 0.0522 | 0.0389 | 0.1613 | 0.8354 | 0.0843 | 0.7941 |
|  | ALL | **0.0021** | 0.0015 | 0.0047 | 0.9978 | 0.0114 | 0.9945 |

ALL: PseAAC+PsePSSM+CT+GO(wMLDAe)

In Table1 that for the virus dataset, when only a single feature extraction method is used, the performance of GO information is the best. The PL index of GO information is 17.31%, 14.50% and 23.71% lower than that of PseAAC, PsePSSM and CT respectively. The values in other indicators in GO information are also significantly better than other single feature extraction algorithms. However, the various indexes of feature fusion data based on wMLDAe dimension reduction show superior characteristics. Its PL, HL, RL, AP, CV and F1 values are 1.33%, 0.64%, 1.44%, 99.61%, 4.66% and 98.85%. Similarly, for the plant dataset, the performance of GO information in the single feature extraction method is the best, which is significantly better than the index values of other feature extraction methods. Its corresponding PL index is 11.38%, 9.32%, and 17.03% lower than PseAAC, PsePSSM, and CT, respectively. And the various indexes of feature fusion data based on wMLDAe dimension reduction show superior characteristics. Its PL, HL, RL, AP, CV, and F1 values are 0.21%, 0.15%, 0.47%, 99.78%, 1.14%, and 99.45%.

Through comparison, this paper chooses the fusion method based on wMLDAe dimension reduction. At this time, the virus dataset obtains a total of 1357-dimensional feature vectors and the plant dataset obtains a total of 3699-dimensional feature vectors. And the dimensions generated by the four feature extraction methods are shown in Table 2.

**Table 2.** Dimensions generated by the four feature extraction methods.

|  | PseAAC | PsePSSM | CT | GO | Total |
| --- | --- | --- | --- | --- | --- |
| Virus | 65D | 200D | 343D | 749D | 1357D |
| Plant | 65D | 200D | 343D | 3091D | 3699D |

### 3.3. Effect of dimensional reduction algorithm on results

In this study, drawing the feature information by four feature extraction algorithms are fused, and many redundant features are also brought. This kind of noise information has some influence on protein SCL. To obtain the optimal feature vector, PCA [46], MVMD [47], MLSI [48], MDDM [49], DMLDA [50] and wMLDAe algorithm are used to reduce the dimensionality. Different dimensionality reduction methods use different dimension combination to obtain different prediction results, so set the dimension to 10 to 100 and the interval to 10. The ProSVM classifier is selected to predict the feature subsets after dimensionality reduction, and the PL is used as the evaluation index. Fig. 5 shows the changes in the PL value of the virus and plant datasets when six different dimensionality reduction methods are selected. When using the wMLDAe algorithm, in order to more clearly analyze the results when the virus and plant datasets take different dimensions, the specific results are shown in Supplementary Table S5 and S6.

**Fig. 5.**
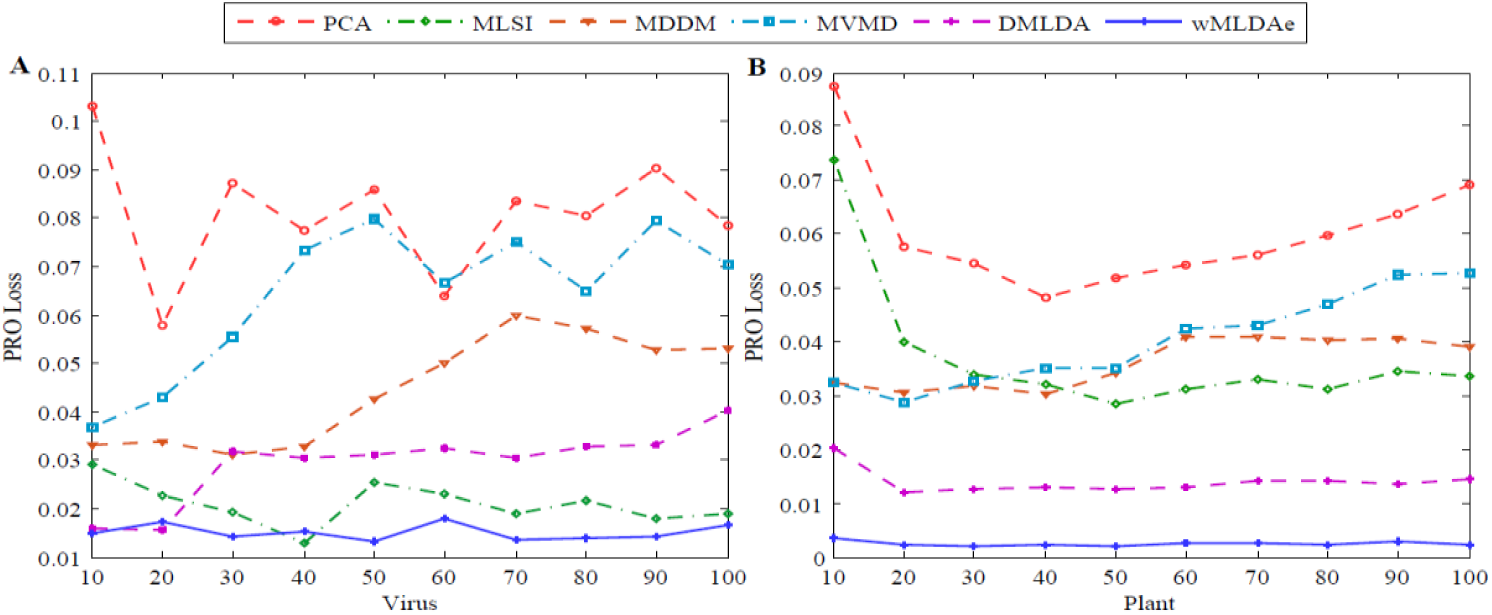
The effect of different dimensionality reduction methods and different dimensions for virus and plant datasets. (A) PL values of virus dataset with different methods and different dimensions (B) PL values of plant dataset with different methods and different dimensions.

It can be known from Fig. 5 that when different dimensionality reduction methods and different dimensions are selected, the prediction results obtained are also different. Compared with other algorithm, wMLDAe algorithm has the best prediction performance. For the virus dataset, when using wMLDAe algorithm to reduce dimensions and selecting the dimension of 50, the lowest PL value is 1.33%, which is 7.27%, 6.63%, 1.21%, 2.93% and 1.79% lower than that of PCA, MVMD, MLSI, MDDM and DMLDA, respectively. For the plant dataset, when using wMLDAe algorithm and selecting 50-dimensions, the PL value is 0.21%, which is 4.97%, 3.29%, 2.63%, 3.21% and 1.05% lower than that of PCA, MVMD, MLSI, MDDM and DMLDA, respectively.

The results show that wMLDAe can reduce noise and redundant information by introducing entropy weight. The wMLDAe algorithm is significantly better than the PCA, MVMD, MLSI, MDDM and DMLDA algorithms. It shows that the algorithm can not only remove the irrelevant information in the feature vector, but also retain the effective features in the sequence and form the optimal feature subset, so as to promote the running speed and increase the prediction accuracy about the model. Therefore, we choose wMLDAe algorithm for dimension reduction and select the dimension of 50. The performance of this model is optimal.

### 3.4. Selection of classification algorithms

The choice of classifier plays a vital role in construction of model. Selecting an appropriate classification algorithm can optimize the complexity of the model and improve the prediction accuracy. For the feature vectors based on wMLDAe dimensionality reduction, this paper uses ML-KNN [38], LIFT [39], Rank-SVM [51], ML-RBF [52], ML-GKR [14] and ProSVM classification algorithms to perform protein SCL prediction on virus and plant datasets, respectively. The jackknife method is used to test, and seven measurement methods such as HL, RL, OAA and OLA are used as evaluation indexes. The prediction results obtained by selecting different classification algorithms for the two training sets are shown in Fig. 6. The specific experimental results for the six classifiers are shown in Supplementary Table S7. For the comparison algorithm, the parameter configuration proposed in this paper is shown in Supplementary Table S8.

**Fig. 6.**
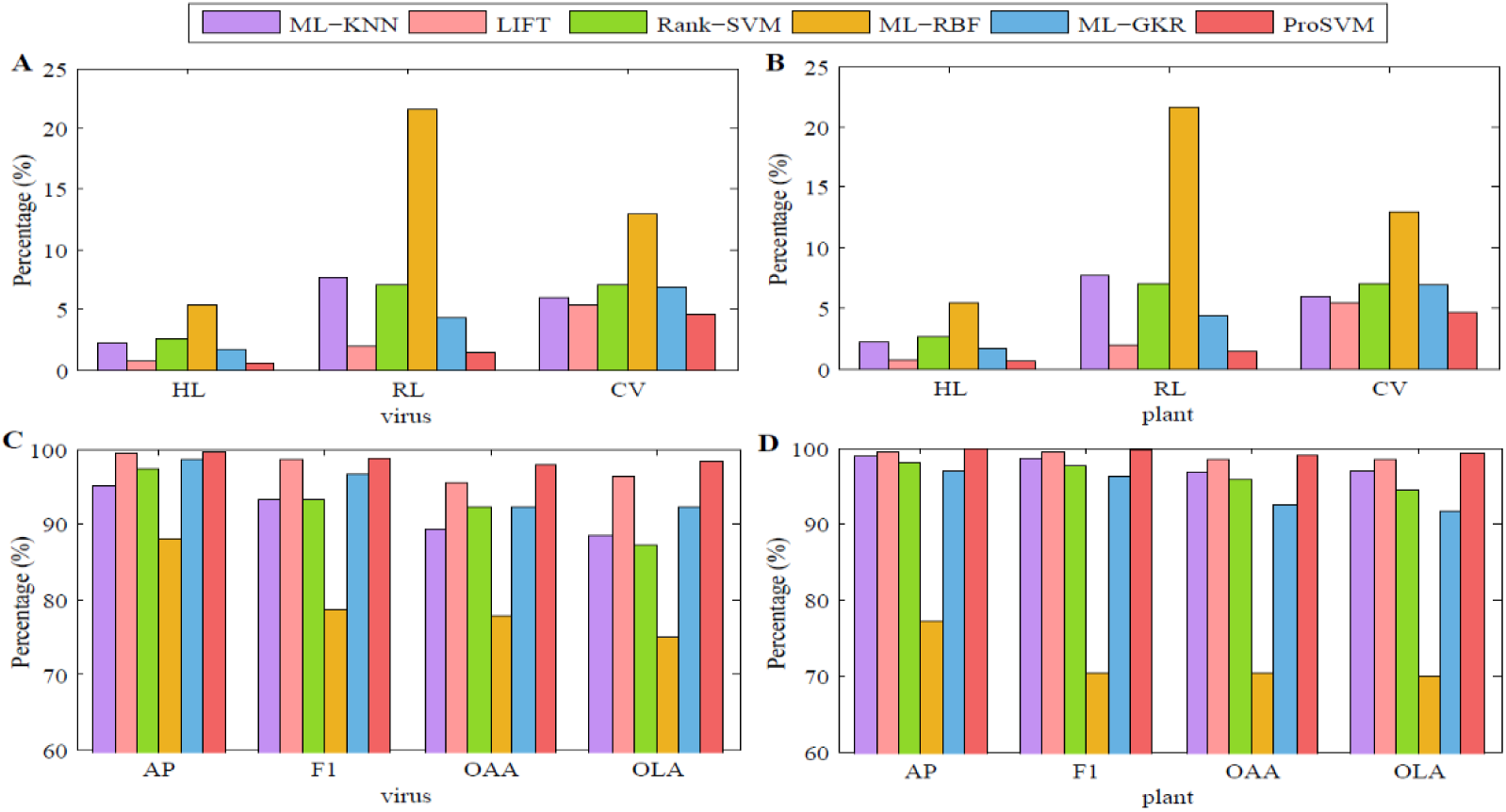
Prediction results for virus and plant datasets when selecting different classifiers. (A) and (C) The results of HL, RL, CV, AP, F1, OAA, OLA values obtained by different classifiers in virus dataset. (B) and (D) The results of HL, RL, CV, AP, F1, OAA, OLA values obtained by different classifiers in plant dataset.

From Fig. 6, when we use the ProSVM algorithm, the prediction performance is the best. For the virus dataset, the OAA and OLA values obtained by using ProSVM algorithm are 98.06% and 98.41% respectively, which are higher than those obtained by using ML-KNN, LIFT, Rank-SVM, ML-RBF and ML-GKR algorithms. The prediction results obtained by using ProSVM algorithm in other indicators are also significantly better than other algorithms. Similarly, for the plant dataset, the OAA and OLA values obtained by ProSVM are 98.97% and 99.33% respectively, which are higher than other algorithms. For other indicators, the prediction results obtained using the ProSVM algorithm are also better than other algorithms.

In Fig. 6, by comparing the prediction results of the virus and plant datasets respectively, we find that the model has the best stability and prediction performance when the ProSVM algorithm is selected. However, the ML-KNN algorithm has the problems of large computational cost and slow prediction speed. For LIFT and Rank-SVM algorithms, both use kernel functions for mapping to high-dimensional space to optimize nonlinear classification problems. Both are susceptible to the choice of kernel functions and parameters, and for Rank-SVM algorithm, it is difficult to run when the amount of data is large. For the ML-RBF algorithm, it is generally combined with K-means clustering, which has the problems of low cluster quality and unstable performance. As the increase of training samples, the number of hidden neurons in RBF network also increases, which increases the complexity of the network and the amount of calculation. From the virus and plant datasets, it can also be concluded that the prediction effect of ML-RBF is not ideal. For ML-GKR algorithm, the benchmark data is decomposed by binary correlation, so that the multi-label problem can be converted into a single-label problem. However, it is necessary to use complete feature information for prediction, and it is easy to lose effectiveness in high-dimensional data. In summary, we select ProSVM algorithm as the classifier to predict the SCL of multi-label proteins.

### 3.5. Comparison with other methods

In the paper, to clarify the prediction performance about this model, we chose to use the same evaluation index for comparison based on the jackknife method. In this paper, the virus and plant datasets are predicted. Firstly, features are extracted and fused by using PseAAC, PsePSSM, GO information and CT, then the fused high-dimensional vector is reduced by using wMLDAe algorithm to get the best feature vector, and finally it is entered into ProSVM to get the prediction results. The comparative results of virus and plant datasets are shown in Table 3 and 4.

**Table 3.** Comparison of prediction results of different methods in virus dataset.

| Locations |  | Methods |  |  |  |  |
| --- | --- | --- | --- | --- | --- | --- |
|  |  | mGOASVM<br>[53] | HybridGO-Loc [54] | mPLR-Loc [55] | MSLVP<br>[12] | ProSVM |
| 1 | Viral capsid | 1.0000 | 1.0000 | 1.0000 | 0.9920 | 1.0000 |
| 2 | Host cell membrane | 0.9700 | 0.9700 | 0.9090 | 0.9920 | 0.9697 |
| 3 | Host endoplasmic<br>reticulum | 0.8500 | 0.9000 | 0.8500 | 0.9960 | 1.0000 |
| 4 | Host cytoplasm | 0.9770 | 0.9660 | 0.9890 | 0.9880 | 0.9885 |
| 5 | Host nucleus | 0.9760 | 0.9880 | 0.964 | 0.9960 | 0.9762 |
| 6 | Secreted | 1.0000 | 1.0000 | 0.8500 | 0.9960 | 1.0000 |
| PL |  | - | - | - | - | 0.0133 |
| HL |  | 0.0260 | 0.0160 | 0.0230 | - | 0.0064 |
| RL |  | - | - | - | - | 0.0144 |
| AP |  | 0.9390 | 0.9650 | 0.957 | - | 0.9961 |
| CV |  | - | - | - | - | 0.0466 |
| F1 |  | 0.9500 | 0.9680 | 0.9550 | - | 0.9885 |
| OAA |  | 0.8890 | 0.9370 | 0.9030 | 0.9750 | **0.9806** |
| OLA |  | 0.9680 | 0.9720 | 0.9480 | 0.9930 | **0.9841** |

*Note:* ‘-’ indicates that the corresponding references do not provide the results of the corresponding indicators.

**Table 4.** Comparison of prediction results of different methods in plant dataset.

| Locations |  | Methods |  |  |  |  |
| --- | --- | --- | --- | --- | --- | --- |
|  |  | iLoc-Plant<br>[56] | mGOASVM<br>[53] | HybridGO-Loc<br>[54] | mPLR-Loc<br>[55] | ProSVM |
| 1 | Cell membrane | 0.6960 | 0.9460 | 0.9110 | 0.8930 | 1.0000 |
| 2 | Cell wall | 0.5940 | 0.8440 | 0.8750 | 0.7810 | 0.9688 |
| 3 | Chloroplast | 0.8810 | 0.9510 | 0.9720 | 0.9830 | 1.0000 |
| 4 | Cytoplasm | 0.6260 | 0.9560 | 0.9230 | 0.9010 | 0.9890 |
| 5 | Endoplasmic<br>reticulum | 0.5000 | 0.9050 | 0.9050 | 0.8330 | 1.0000 |
| 6 | Extracellular | 0.0910 | 1.0000 | 0.9550 | 0.8640 | 1.0000 |
| 7 | Golgi apparatus | 0.7620 | 0.9050 | 0.9050 | 0.8570 | 0.9524 |
| 8 | Mitochondrion | 0.7470 | 1.0000 | 0.9930 | 0.9930 | 1.0000 |
| 9 | Nucleus | 0.9210 | 0.9930 | 0.9870 | 0.9610 | 0.9934 |
| 10 | Peroxisome | 0.2860 | 1.0000 | 1.0000 | 1.0000 | 1.0000 |
| 11 | Plastid | 0.1790 | 1.0000 | 0.9740 | 0.9230 | 0.9744 |
| 12 | Vacuole | 0.5380 | 0.9420 | 0.9230 | 0.9420 | 0.9808 |
| PL |  | - | - | - | - | 0.0021 |
| HL |  | - | 0.0130 | 0.0070 | 0.0100 | 0.0015 |
| RL |  | - | - | - | - | 0.0047 |
| AP |  | - | 0.9330 | 0.9720 | 0.9560 | 0.9978 |
| CV |  | - | - | - | - | 0.0114 |
| F1 |  | - | 0.9420 | 0.9660 | 0.9490 | 0.9945 |
| OAA |  | 0.6810 | 0.8740 | 0.9360 | 0.9080 | **0.9897** |
| OLA |  | 0.7170 | 0.9620 | 0.9560 | 0.9370 | **0.9933** |

Note: '-' indicates that the corresponding references do not provide the results of the corresponding indicators.

It can be seen from Table 3 and 4, for virus dataset, when using MpsLDA-ProSVM method, the OAA and OLA are 98.06% and 98.41% respectively, which are 4.36%∼9.16% and 1.21%∼3.61% higher than those using mGOASVM, HybridGO-Loc and mPLR-Loc methods. Although the OLA value obtained by using the MSLVP method is 0.89% higher than that under the MpsLDA-ProSVM method, the OAA value is 0.56% lower than that. For plant dataset, when using MpsLDA-ProSVM method, the OAA and OLA are 98.97% and 99.33% respectively, which are 5.37%∼30.87% and 3.13%∼27.63% higher than those using iLoc-Plant, mGOASVM, HybridGO-Loc and mPLR-Loc methods. Moreover, MpsLDA-ProSVM is superior to other methods in the prediction of each location of the two datasets.

So as to further verify the rationality and validity about this model, the datasets of Gram-positive bacteria and Gram-negative bacteria are used as test sets, and by the jackknife method, we obtain the specific prediction results about two test sets. Selecting the same dataset for comparison, the comparison results are shown in Table 5 and Table 6. For Gram-positive bacteria dataset, when using MpsLDA-ProSVM method, the OAA and OLA are all 99.81%, which are 3.51%∼6.91% and 3.01%∼6.71% higher than those using other methods, respectively. For Gram-negative bacteria dataset, when using MpsLDA-ProSVM method, the OAA and OLA are 98.49% and 99.04% respectively, which are 3.99%∼8.59% and 3.74%∼7.64% higher than those using other methods. And in each protein, the prediction accuracy obtained by using MpsLDA-ProSVM method is obviously better than other methods. Therefore, the prediction model proposed in this paper has good generalization ability and achieves satisfactory results.

**Table 5.**
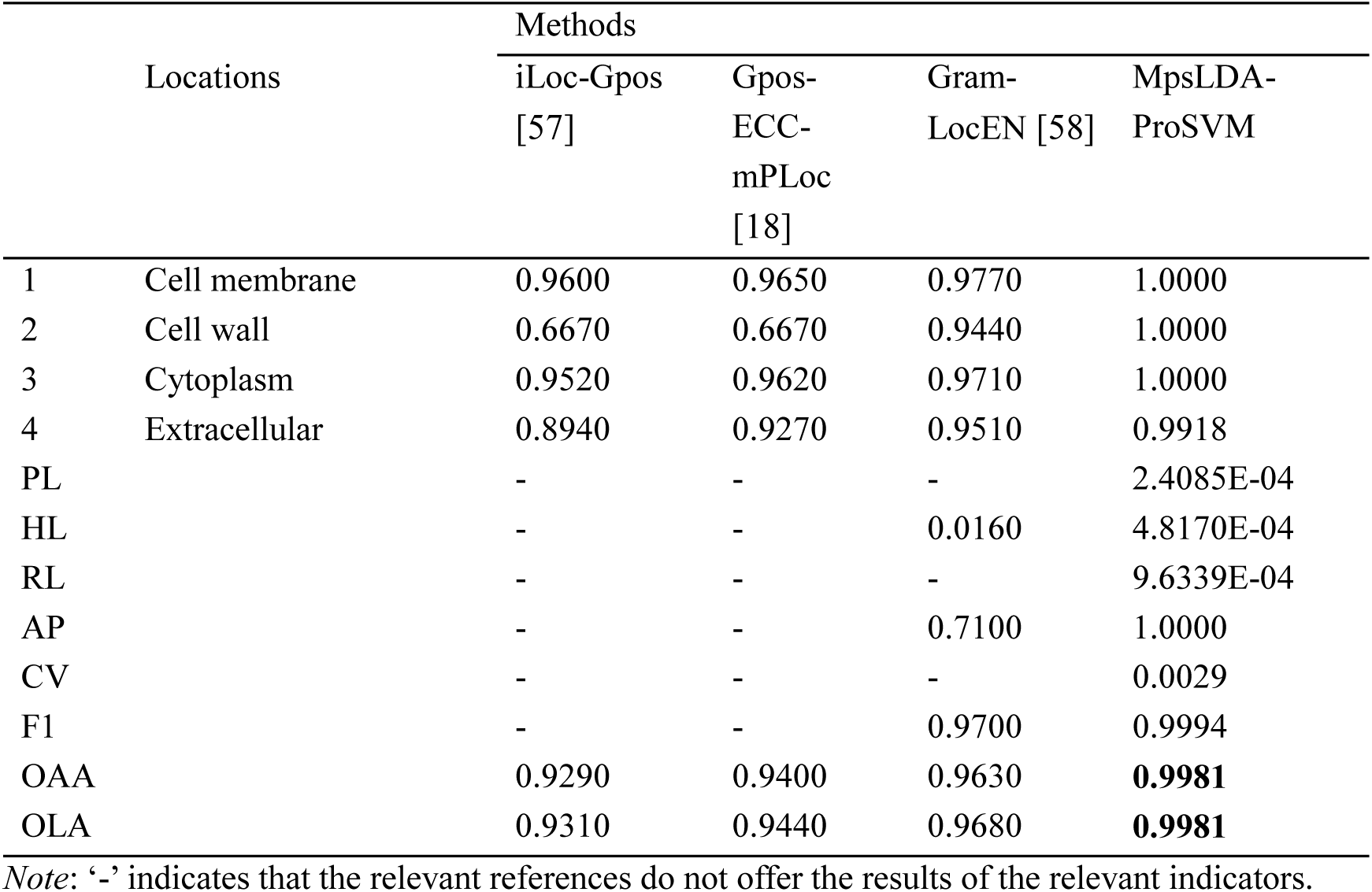
Comparison of different prediction methods for protein SCL on Gram-positive bacteria.

**Table 6.**
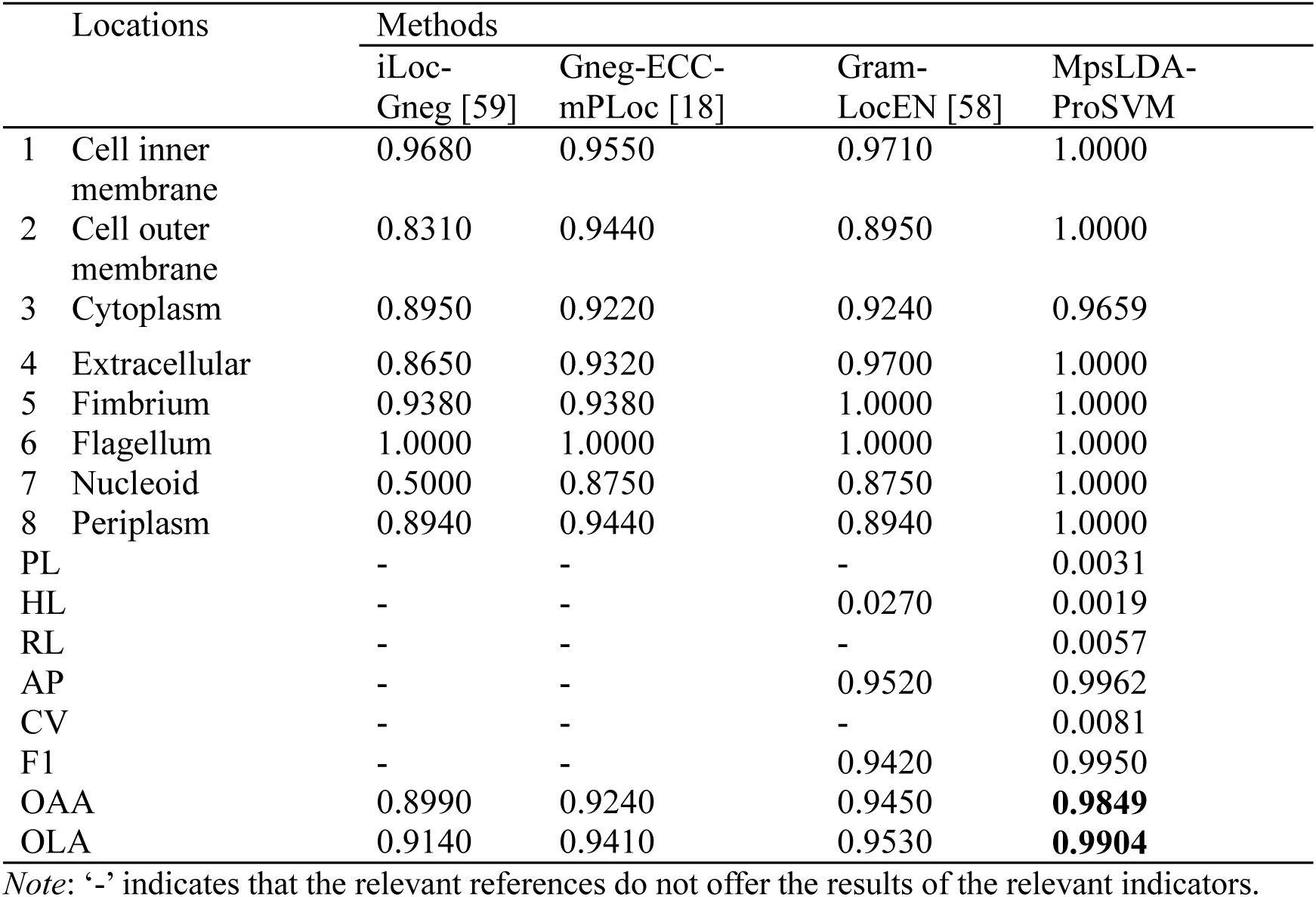
Comparison of different prediction methods for protein SCL on Gram-negative bacteria.

## 4. Conclusion

The precise location of the SCL of multi-label proteins can not only clarify the various functions of proteins, but also promote the knowledge of the pathogenesis with some diseases and the research of new drugs. This paper proposes a new method, MpsLDA-ProSVM, to predict the SCL of multi-label proteins. First, the fusion of the feature vectors extracted by PseAAC, PsePSSM, GO and CT can fully mine the annotation information, sequence information, evolution information and physicochemical information in protein sequences. Secondly, in order to avoid dimension disaster, wMLDAe is used for the first time to remove irrelevant information. It applies the entropy weight form to the MLDA method, fully considers the label information, and improves the speed of the model. Finally, the feature vector is input to the ProSVM classifier to predict multi-label proteins. For getting the comprehensive ranking of relevant labels, we first use ML-KNN and LIFT algorithm to get the predicted label scores of each instance, summarize the scores and sort them to get the ranking, and then the objective function is optimized according to the ADMM algorithm to obtain the final prediction result. This algorithm further improves the efficiency of the model. Combined with the above methods, the model has achieved satisfactory prediction results and is obviously superior to other prediction methods. With the continuous collection and identification of multi-label proteins, we expect that MpsLDA-ProSVM can be used on datasets of more species. In recent years, deep learning has successfully predicted the sites of multi-label data by virtue of its powerful generalization ability. In the future, we will try to use the deep learning to predict multi-label protein subcellular location.

## Declaration of Competing Interest

The authors declare that they have no known competing financial interests or personal relationships that could have appeared to influence the work reported in this paper.

## Acknowledgments

This work was supported by the National Nature Science Foundation of China (61863010, 61877064, U1806202), the Key Research and Development Program of Shandong Province of China (2019GGX101001), and the Natural Science Foundation of Shandong Province of China (ZR2018MC007).

## Notes

### Competing Interest Statement

The authors have declared no competing interest.

