## Supplementary Table S1-Table S8 for "MpsLDA-ProSVM: predicting multi-label protein subcellular localization by wMLDAe dimensionality reduction and ProSVM classifier"

### Table of Contents

#### 1. Supplementary Tables

Table S1. Breakdown of the Gram-positive bacterial benchmark dataset.

Table S2. Breakdown of the Gram-negative bacterial benchmark dataset

Table S3. Select different parameter $\lambda$ to obtain prediction results of virus and plant datasets..

Table S4. Select different parameter $\xi$ to obtain prediction results of virus and plant datasets.

Table S5. Prediction results obtained by selecting different dimensions when the virus dataset uses wMLDAe for dimensionality reduction.

Table S6. Prediction results obtained by selecting different dimensions when the plant dataset uses wMLDAe for dimensionality reduction.

Table S7. Comparison of the results of different classification algorithms for virus and plant datasets.

Table S8. Parameter configuration of six algorithms.

#### 1. Supplementary Tables

**Table S1.** Breakdown of the Gram-positive bacterial benchmark dataset. None of the positive proteins included here has ≥ 25% sequence identity to any other in a same subcellular location.

| Subset | Subcellular location | Number of proteins |
|---|---|---|
| S1 | Cell membrane | 174 |
| S2 | Cell wall | 18 |
| S3 | Cytoplasm | 208 |
| S4 | Extracellular | 123 |
| Total number of locative proteins | | 523 |
| Total number of different proteins | | 519 |

**Table S2.** Breakdown of the Gram-negative bacterial benchmark dataset. None of the negative proteins included here has ≥ 25% sequence identity to any other in a same subcellular location.

| Subset | Subcellular location | Number of proteins |
|---|---|---|
| S1 | Cell inner membrane | 557 |
| S2 | Cell outer membrane | 124 |
| S3 | Cytoplasm | 410 |
| S4 | Extracellular | 133 |
| S5 | Fimbrium | 32 |
| S6 | Flagellum | 12 |
| S7 | Nucleoid | 8 |
| S8 | Periplasm | 180 |
| Total number of locative proteins | | 1456 |
| Total number of different proteins | | 1392 |

**Table S3.** Select different parameter $\lambda$ to obtain prediction results of virus and plant datasets.

| Dataset | Parameter | PL | HL | RL | AP | CV | F1 |
|---|---|---|---|---|---|---|---|
| | 5 | 0.2183 | 0.2520 | 0.2184 | O.6759 | 0.3178 | 0.4684 |
| | 10 | 0.1972 | 0.2383 | 0.1942 | 0.6985 | 0.3064 | 0.4956 |
| | 15 | 0.1984 | 0.2423 | 0.1921 | 0.7025 | 0.3033 | 0.5004 |
| | 20 | 0.1948 | 0.2391 | 0.1978 | 0.6919 | 0.2973 | 0.5083 |
| | 25 | 0.1895 | 0.2334 | 0.1854 | 0.7115 | 0.2845 | 0.5157 |
| Virus | 30 | 0.1920 | 0.2318 | 0.1847 | 0.7102 | 0.2861 | 0.5054 |
| | 35 | 0.1862 | 0.2214 | 0.1768 | 0.7149 | 0.2790 | 0.5280 |
| | 40 | 0.1868 | 0.2262 | 0.1778 | 0.7243 | 0.2869 | 0.5338 |
| | 45 | 0.1864 | 0.2326 | 0.1741 | 0.7270 | 0.2842 | 0.5146 |
| | 49 | 0.1852 | 0.2326 | 0.1819 | 0.7125 | 0.2913 | 0.194 |
| | 5 | 0.1795 | 0.1910 | 0.1710 | 0.5909 | 0.0963 | 0.3891 |
| | 10 | 0.1737 | 0.2052 | 0.1735 | 0.5815 | 0.0858 | 0.3884 |

| | | | | | | | |
|---|---|---|---|---|---|---|---|
| | 15 | 0.1740 | 0.2131 | 0.1757 | 0.5730 | 0.0913 | 0.3799 |
| | 20 | 0.1738 | 0.2227 | 0.1749 | 0.5762 | 0.0869 | 0.3776 |
| | 25 | 0.1698 | 0.2256 | 0.1753 | 0.5746 | 0.0895 | 0.3822 |
| Plant | 30 | 0.1715 | 0.2296 | 0.1779 | 0.5698 | 0.0872 | 0.3766 |
| | 35 | 0.1706 | 0.2321 | 0.1440 | 0.5817 | 0.0857 | 0.3729 |
| | 40 | 0.1679 | 0.2286 | 0.1757 | 0.5699 | 0.0839 | 0.3810 |
| | 45 | 0.1660 | 0.2292 | 0.1717 | 0.5815 | 0.0841 | 0.3861 |
| | 49 | 0.1704 | 0.2349 | 0.1730 | 0.5809 | 0.0873 | 0.3733 |

**Table S4.** Select different parameter $\xi$ to obtain prediction results of virus and plant datasets.

| Dataset | Parameter | PL | HL | RL | AP | CV | F1 |
|---|---|---|---|---|---|---|---|
| | 1 | 0.1694 | 0.2648 | 0.1774 | 0.7440 | 0.2594 | 0.5449 |
| | 2 | 0.1663 | 0.2640 | 0.1731 | 0.7489 | 0.2536 | 0.5492 |
| | 3 | 0.1646 | 0.2520 | 0.1699 | 0.7517 | 0.2420 | 0.5577 |
| | 4 | 0.1653 | 0.2592 | 0.1647 | 0.7600 | 0.2572 | 0.5538 |
| | 5 | 0.1681 | 0.2544 | 0.1700 | 0.7446 | 0.2548 | 0.5491 |
| Virus | 6 | 0.1675 | 0.2536 | 0.1662 | 0.7549 | 0.2495 | 0.5468 |
| | 7 | 0.1620 | 0.2520 | 0.1647 | 0.7620 | 0.2471 | 0.5631 |
| | 8 | 0.1647 | 0.2592 | 0.1649 | 0.7596 | 0.2330 | 0.5477 |
| | 9 | 0.1583 | 0.2512 | 0.1630 | 0.7574 | 0.2439 | 0.5607 |
| | 10 | 0.1646 | 0.2608 | 0.1691 | 0.7532 | 0.2447 | 0.5484 |
| | 1 | 0.1490 | 0.2258 | 0.1621 | 0.6014 | 0.0872 | 0.4004 |
| | 2 | 0.1496 | 0.2263 | 0.1617 | 0.6068 | 0.0907 | 0.4012 |
| | 3 | 0.1496 | 0.2257 | 0.1597 | 0.6075 | 0.0930 | 0.4017 |
| | 4 | 0.1490 | 0.2238 | 0.1586 | 0.6072 | 0.0917 | 0.4045 |
| | 5 | 0.1474 | 0.2205 | 0.1573 | 0.6119 | 0.0892 | 0.4090 |
| Plant | 6 | 0.1469 | 0.2206 | 0.1578 | 0.6111 | 0.0883 | 0.4085 |
| | 7 | 0.1521 | 0.2289 | 0.1698 | 0.5917 | 0.0873 | 0.3974 |
| | 8 | 0.1458 | 0.2167 | 0.1575 | 0.6121 | 0.0885 | 0.4155 |
| | 9 | 0.1454 | 0.2164 | 0.1559 | 0.6139 | 0.0859 | 0.4159 |
| | 10 | 0.1466 | 0.2156 | 0.1557 | 0.6128 | 0.0814 | 0.4156 |

It can be seen from Table S3 and Table S4 that the prediction result of the classifier varies with different parameter values. The smaller the PL value, the better the performance of the model. For the value of parameter $\lambda$, when the parameter value of virus dataset is 49, the PL value is the lowest, 3.31% lower than the initial value of $\lambda$=5. When the parameter value of plant dataset is 45, the PL value is the lowest, which is 1.35% lower than the initial value $\lambda$=5. The PL value of plant dataset is 0.44% higher than that of $\lambda$=45 when $\lambda$=49, and that of virus dataset is 0.12% higher than that of $\lambda$=49 when $\lambda$=45. Therefore, $\lambda$=45 is selected under comprehensive consideration. At this time, when using PseAAC for feature extraction, a total of (20+$\lambda$=65)

-dimensional feature vectors are generated. For the value of parameter $\xi$, when the parameter value of the virus dataset is 9, the PL is the smallest, which is 1.11% lower than the PL value when the initial value is $\xi=1$. When the parameter value of plant dataset is 9, the PL value is the lowest, 0.67% lower than that of the highest $\xi=7$. Therefore, when using PsePSSM algorithm for feature extraction, select $\xi=9$. At this time, a protein sequence generates a $(20+20\times\xi=200)$ -dimensional feature vector.

**Table S5.** Prediction results obtained by selecting different dimensions when the virus dataset uses wMLDAe for dimensionality reduction. (Where ↓ means the smaller the index value, the better; ↑ means the larger the index value, the better)

| Metric | Dimension | | | | | | | | | |
|---|---|---|---|---|---|---|---|---|---|---|
| | Jackknife test | | | | | | | | | |
| | 10 | 20 | 30 | 40 | 50 | 60 | 70 | 80 | 90 | 100 |
| PL↓ | 0.0149 | 0.0173 | 0.0142 | 0.0152 | **0.0133** | 0.0179 | 0.0136 | 0.0138 | 0.0142 | 0.0165 |
| HL↓ | 0.0072 | 0.0080 | 0.0048 | 0.0048 | 0.0064 | 0.0088 | 0.0048 | 0.0048 | 0.0032 | 0.0048 |
| RL↓ | 0.0171 | 0.0193 | 0.0112 | 0.0104 | 0.0144 | 0.0225 | 0.0120 | 0.0112 | 0.0072 | 0.0104 |
| AP↑ | 0.9988 | 0.951 | 0.9972 | 0.9961 | 0.9961 | 0.9937 | 0.9969 | 0.9959 | 0.9981 | 0.9961 |
| CV↓ | 0.0556 | 0.0515 | 0.0475 | 0.0426 | 0.0466 | 0.0547 | 0.0483 | 0.0442 | 0.0434 | 0.0426 |
| F1↑ | 0.9871 | 0.9863 | 0.9924 | 0.9925 | 0.9885 | 0.9819 | 0.9922 | 0.9924 | 0.9954 | 0.9925 |
| OAA ↑ | 0.9565 | 0.9661 | 0.9661 | 0.9468 | **0.9806** | 0.9516 | 0.9710 | 0.9613 | 0.9613 | 0.9516 |
| OLA↑ | 0.9563 | 0.9682 | 0.9682 | 0.9404 | 0.9841 | 0.9563 | 0.9603 | 0.9563 | 0.9603 | 0.9484 |

**Table S6.** Prediction results obtained by selecting different dimensions when the plant dataset uses wMLDAe for dimensionality reduction.

| Metric | Dimension | | | | | | | | | |
|---|---|---|---|---|---|---|---|---|---|---|
| | Jackknife test | | | | | | | | | |
| | 10 | 20 | 30 | 40 | 50 | 60 | 70 | 80 | 90 | 100 |
| PL↓ | 0.0035 | 0.0024 | 0.0022 | 0.0024 | **0.0021** | 0.0025 | 0.0027 | 0.0022 | 0.0028 | 0.0022 |
| HL↓ | 0.0049 | 0.0012 | 0.0011 | 0.0016 | 0.0015 | 0.0012 | 0.0017 | 0.0012 | 0.0010 | 0.0010 |
| RL↓ | 0.0079 | 0.0037 | 0.0032 | 0.0044 | 0.0047 | 0.0037 | 0.0045 | 0.0041 | 0.0028 | 0.0027 |
| AP↑ | 09882 | 0.9985 | 0.9988 | 0.9983 | 0.9978 | 0.9985 | 0.9982 | 0.9985 | 0.9981 | 0.9978 |
| CV↓ | 0.0124 | 0.0108 | 0.0102 | 0.0112 | 0.0114 | 0.0108 | 0.0111 | 0.0116 | 0.0094 | 0.0092 |
| F1↑ | 0.9812 | 0.9958 | 0.9962 | 0.9942 | 0.9945 | 0.9958 | 0.9938 | 0.9958 | 0.9963 | 0.9961 |
| OAA ↑ | 0.9519 | 0.9867 | 0.9887 | 0.9867 | **0.9897** | 0.9867 | 0.9887 | 0.9887 | **0.9897** | 0.9887 |
| OLA↑ | 0.9924 | 0.9914 | 0.9933 | 0.9914 | 0.9933 | 0.9905 | 0.9924 | 0.9933 | 0.9943 | 0.9924 |

**Table S7.** Comparison of the results of different classification algorithms for virus and plant datasets.

| Dataset | Algorithm | HL | RL | AP | CV | F1 | OAA | OLA |
|---|---|---|---|---|---|---|---|---|
| Virus | MLKNN | 0.0225 | 0.0775 | 0.9507 | 0.0595 | 0.9318 | 0.8937 | 0.8849 |
| | LIFT | 0.0072 | 0.0191 | 0.9942 | 0.0539 | 0.9869 | 0.9565 | 0.9642 |
| | MLSVM | 0.0257 | 0.0700 | 0.9734 | 0.0700 | 0.9323 | 0.9227 | 0.8730 |
| | MLRBF | 0.0539 | 0.2159 | 0.8800 | 0.1296 | 0.7864 | 0.7778 | 0.7500 |
| | MLGKR | 0.0161 | 0.0432 | 0.9858 | 0.0684 | 0.9674 | 0.9227 | 0.9246 |
| | ProSVM | 0.0064 | 0.0144 | 0.9961 | 0.0466 | 0.9885 | **0.9806** | 0.9841 |
| Plant | MLKNN | 0.0035 | 0.0163 | 0.9879 | 0.0197 | 0.9868 | 0.9662 | 0.9687 |
| | LIFT | 0.0015 | 0.0075 | 0.9947 | 0.0120 | 0.9943 | 0.9836 | 0.9848 |
| | MLSVM | 0.0051 | 0.0291 | 0.9803 | 0.0243 | 0.9751 | 0.9580 | 0.9431 |
| | MLRBF | 0.0277 | 0.2960 | 0.7704 | 0.1315 | 0.7041 | 0.7034 | 0.6995 |
| | MLGKR | 0.0077 | 0.0465 | 0.9695 | 0.0437 | 0.9630 | 0.9253 | 0.9156 |
| | ProSVM | 0.0015 | 0.0047 | 0.9978 | 0.0114 | 0.9945 | **0.9897** | 0.9933 |

For virus dataset, the OAA and OLA values obtained by using ProSVM algorithm are 98.06% and 98.41% respectively, which are 8.69% and 9.92%, 2.41% and 1.99%, 5.79% and 11.11%, 20.28% and 23.41%, 5.79% and 5.95% higher than those obtained by using ML-KNN, LIFT, Rank-SVM, ML-RBF and ML-GKR algorithms. The prediction results obtained by using ProSVM algorithm in other indicators are also significantly better than other algorithms. For plant dataset, the OAA and OLA values obtained by ProSVM are 98.97% and 99.33% respectively, which are 2.35% and 2.46%, 0.61% and 0.85%, 3.17% and 5.02%, 28.63% and 29.38%, 6.44% and 7.77% higher than those obtained by ML-KNN, LIFT, Rank-SVM, ML-RBF and ML-GKR algorithms. For other indicators, the prediction results obtained using the ProSVM algorithm are also better than other algorithms. By comparing the prediction results of the virus and plant datasets respectively, we find that the model has the best stability and prediction performance when the ProSVM algorithm is selected.

**Table S8.** Parameter configuration of six algorithms.

| Algorithm | Parameter |
|---|---|
| ProSVM | $\lambda = 4$, $str\_tag = ProSVM$, $part\_size = 10^7$ |
| ML-KNN | $num = 9$, $smooth = 1$ |
| LIFT | $svm.type = linear$, $svm.para = []$, $ratio = 0.1$ |

| | |
|---|---|
| Rank-SVM | $svm.type = linear$ , $svm.para = []$ , $C = 1$, $\lambda$=$10^{-6}$ , $\alpha = 10^{-4}$ , $max\_iter = 50$ |
| ML-RBF | $ratio = 0.1$, $mu = 0.05$ |
| ML-GKR | $para = 10$ |
